# *Candidatus* Liberibacter asiaticus reduces callose and reactive oxygen species production in the phloem

**DOI:** 10.1101/2022.03.10.483847

**Authors:** Chiara Bernardini, Donielle Turner, Chunxia Wang, Stacy Welker, Diann Achor, Yosvanis Acanda Artiga, Robert Turgeon, Amit Levy

## Abstract

Huanglongbing (HLB) causes significant economic loss in citrus production worldwide. HLB is caused by *Candidatus* Liberibacter asiaticus (*C*Las), a gram-negative bacterium which inhabits the phloem exclusively. *C*Las infection results in accumulation of callose and reactive oxygen species in the phloem of infected plants, but little is known about the specific processes that take place during infection because of the sparse distribution of bacteria and the inaccessibility of the phloem inside the tree. In this study, we used the seed vasculatures, which accumulate a high number of *C*Las, as a model tissue to study *C*Las-host cellular interactions. In vasculature where *C*Las is abundant, sieve pore callose and H_2_O_2_ concentration were reduced compared to healthy seed vasculature. The expression of callose synthases (*CalS*) and respiratory burst oxidase homolog (*RBOH*) genes were downregulated in infected seeds compared to healthy ones. In leaves of HLB-infected plants, H_2_O_2_ concentration and *CalS* expression increased compared to uninfected leaves, but cells with *C*Las had lower levels of sieve plate callose compared to cells without *C*Las. Our results provide evidence that the bacteria manipulate cell metabolism to disable plant defenses and suggests that HLB disease is the result of a constant arms-race between the pathogen and a defense response, which is ultimately harmful to the host plant.

## Introduction

Huanglongbing (HLB) is a severe obstacle for the growth and production of citrus trees. In Florida, HLB has reduced the production of citrus by 74% (Graham et al., 2020). The HLB putative agent is *Candidatus* Liberibacter asiaticus (*C*Las), a gram-negative bacterium which is an obligate inhabitant of the phloem. *C*Las is transmitted by the vector insect *Diaphorina citri* (Kuwaykama) (Galdeano et al., 2020; Hall et al., 2013). The efficiency of transmission by the vector together with the host-pathogen coevolution has resulted in worldwide spread of the disease. In Florida, HLB affects 95% of citrus trees (Kramer et al., 2020). HLB causes a wide variety of symptoms such as yellow shoots, leaves with blotchy yellow mottling (Bové, 2006), loss of fruit (Dala-Paula et al., 2019), and decrease in the quality of fruit production (Chin et al., 2014; Ferguson et al., 2021; Martinelli and Dandekar, 2017). Tree death occurs within months to years after infection (Dala-Paula et al., 2019).

After infection, the concentrations of some regulatory amino, organic, and fatty acids in the shoot are unbalanced (Killiny and Nehela, 2017), and several metabolic pathways are abnormally upregulated or downregulated. Specifically, the plant slows down pathways related to cell wall development and lipid and nucleotide metabolism (Kim et al., 2009), and speeds up phytohormone and pathogenesis-related (PR) protein defense pathways (Hu et al., 2017; Nehela et al., 2018). The pathogen elicits the disruption of sugar transport genes (Kim et al., 2009), resulting in a carbohydrate imbalance which occurs in infected asymptomatic and symptomatic plants (Fan et al., 2010). This interference with basic metabolic functions may be the reason for the quality loss in the fruit (Chin et al., 2014). The affected pathways are modulated at different stages of infection (Folimonova and Achor, 2010; Zheng and Zhao, 2013). Infected plants also show modifications of the phloem tissue, where plasma membrane malformations and deposition of phloem protein 2 (PP2) occur (Achor et al., 2010). This results in the loss of functionality of the phloem and detrimental cytologic modifications (Folimonova and Achor, 2010).

Callose is a polymer of β-1,3 glucan that forms structural components of plant cells (Granato et al., 2019). Callose synthesis is catalyzed by a multi-subunit enzyme complex (Verma and Hong, 2001) that includes sucrose synthase, UDP-glucose transferase, and callose synthase. Callose synthases (CalS) are transmembrane proteins (Nedukha, 2015) responsible for the callose assembly starting from UDP-glucose (Verma and Hong, 2001). In *Arabidopsis*, 12 CALS have been reported to take part in various processes (Ellinger and Voigt, 2014), but only CALS7 is specifically responsible for the callose deposition around the sieve pore (Barratt et al., 2011; Xie et al., 2011). In HLB-infected citrus, *Cals2, Cals7*, and *Cals12* are significantly upregulated (Granato et al., 2019). Moreover, previous work shows that *C*Las infection results in constriction of the sieve pores due to callose deposition (Achor et al., 2010). It has been hypothesized that the plant attempts to plug the sieve elements with callose or PP2 to limit the spread of the pathogen and its effectors (Achor et al., 2010; Deng et al., 2019; Kim et al., 2009; Koh et al., 2012; Welker et al., 2021). Indeed, a complete occlusion of the sieve pores results in the loss of the functionality of the entire vessel (Etxeberria et al., 2009). Without normal sugar translocation, starch accumulates inside the thylakoid system, causing its breakdown, resulting in the loss of functionality of the photosynthetic apparatus (Etxeberria et al., 2009; Granato et al., 2019). In support of the hypothesis that HLB causes photosynthate transport disruption, it has been demonstrated that infected trees have a decreased speed of sugar export and translocation (Koh et al., 2012; Welker et al., 2021).

Another plant response against phloem-restricted pathogens is programmed cell death (Torres et al., 2006). Cell death can be caused by an increase of reactive oxygen species (ROS) inside the cells (Gechev et al., 2006). The increased ROS leads to thickening of the cell wall, damage to the cell membrane and the peroxidation of lipids, culminating in cell death (Torres, 2006). During *C*Las invasion, the production of H_2_O_2_ increases in the leaves, causing a toxic build-up in the tissue and leading to the death of the cells in the surrounding tissue (Pitino et al., 2017). The O_2_^-^ ions, which are the substrate for the peroxidase enzyme family, are provided *in planta* by the respiratory burst oxidase homologs (RBOH) (Ogasawara et al., 2008; Torres et al., 2002). RBOHs have been shown to play an active role during infection (Yoshioka et al., 2003; Yu et al., 2020). In potato, an RBOH protein, among others, was found inside the phloem (Otulak-Kozieł et al., 2019). To control ROS homeostasis, plants employ a scavenger mechanism which is responsible for balancing their concentration inside the cells and to avoid the toxicity for the plant (Shigeoka et al., 2002). The H_2_O_2_ scavengers are either enzymatic, such as catalase (CAT), superoxide dismutase (SOD), ascorbate peroxidase (APX), and gluconate peroxidase (GPX), or low molecular mass antioxidants, which are non-enzymatic (Huang et al., 2019). During *C*Las infection, the plant may attempt to protect the tissue from oxidative stress with proline (Killiny and Nehela, 2017).

Our understanding of *C*Las-plant interactions has been limited due to the low levels of the bacteria in the plant phloem (Louzada et al., 2016). The cultivation of *C*Las in artificial media is difficult (Sechler et al., 2009). Moreover, qPCR and TEM studies show that the bacteria are not equally distributed in all the cells of an infected plant (Achor et al., 2020; Louzada et al., 2016). However, previous studies have shown that large amounts of viable *C*Las cells accumulate in seed phloem of infected plants (Achor et al., 2020; Hilf et al., 2013).

Here, we studied the cellular interaction of *C*Las with the host cell by exploring the vasculatures of healthy and *C*Las-infected seeds and leaves with a combined molecular and microscopy approach. Our results show that in the sieve elements which contained the bacteria, there was no accumulation of callose and a weaker ROS response to the pathogen (H_2_O_2_ in particular) compared to uninfected phloem. In the leaves, sieve element cells containing *C*Las showed less constriction of the sieve pores. Moreover, citrus *CalS* (*CsCalS*) and *RBOH* (*CsRBOH*) genes are strongly downregulated in *C*Las-containing cells compared to healthy ones. These results provide novel evidence that *C*Las can locally inhibit two of the most important plant responses to infection. This inhibition appears to enable direct cell-to-cell movement so that the bacteria can bypass plant defenses locally, thus allowing them to colonize the entire plant.

## RESULTS

### In seed vasculatures, *C*Las is numerous and callose is scarce

In both sweet orange and grapefruit seed vasculatures, high numbers of bacterial cells were observed inside sieve elements (SE) using TEM (Fig. 1A-B). Morphology of the bacteria varied greatly, with tubular, spherical, or flask-shaped forms. The fluorescent in situ hybridization (FISH) analysis confirmed the presence of *C*Las in grapefruit seed vasculature (Fig. 1C-D). It was previously shown that HLB infection causes callose accumulation in leaves and stems, but in seed vasculatures where *C*Las was physically present, sieve plate pores were not occluded by callose (Achor et al., 2020). To determine whether this resulted from a lack of callose deposition in the seeds, or from the presence of *C*Las, we compared infected seed vasculature to uninfected ones. Thick deposits of callose were present at the sieve plates of healthy sweet orange and grapefruit seed vasculature (Fig. 2A and C).

**Figure 1.**
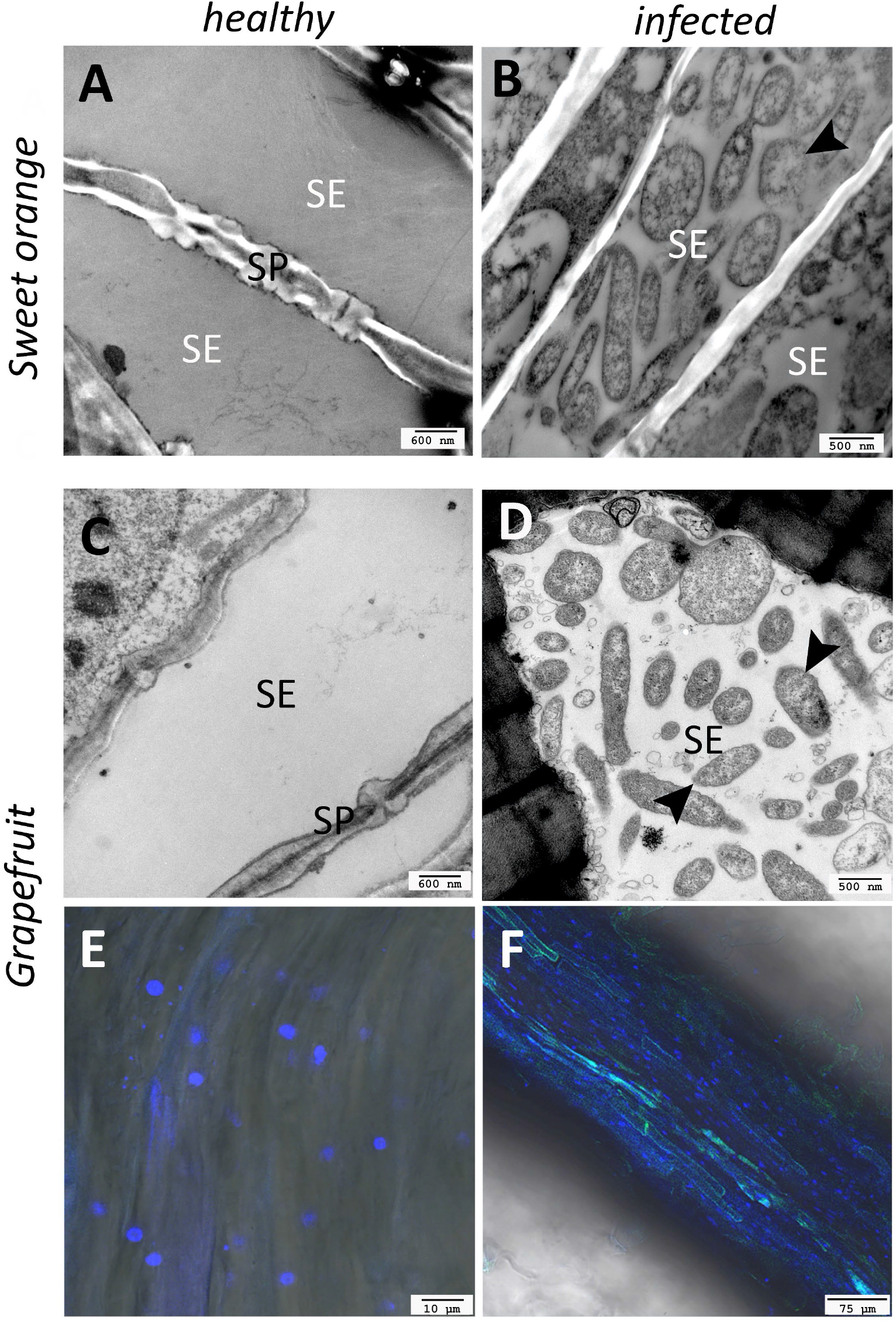
*C*Las accumulates in the seed coat vasculature. (A-D) TEM micrographs of seed vasculature of ‘Valencia’ Sweet Orange (C) and ‘Duncan’ grapefruit (D). (E-F) FISH (fluorescent in situ hybridization) micrographs of healthy and infected seed vasculatures of ‘Duncan’ Grapefruit. Labels: SE= sieve element, SP=sieve plates, triangles= CLas.

**Figure 2.**
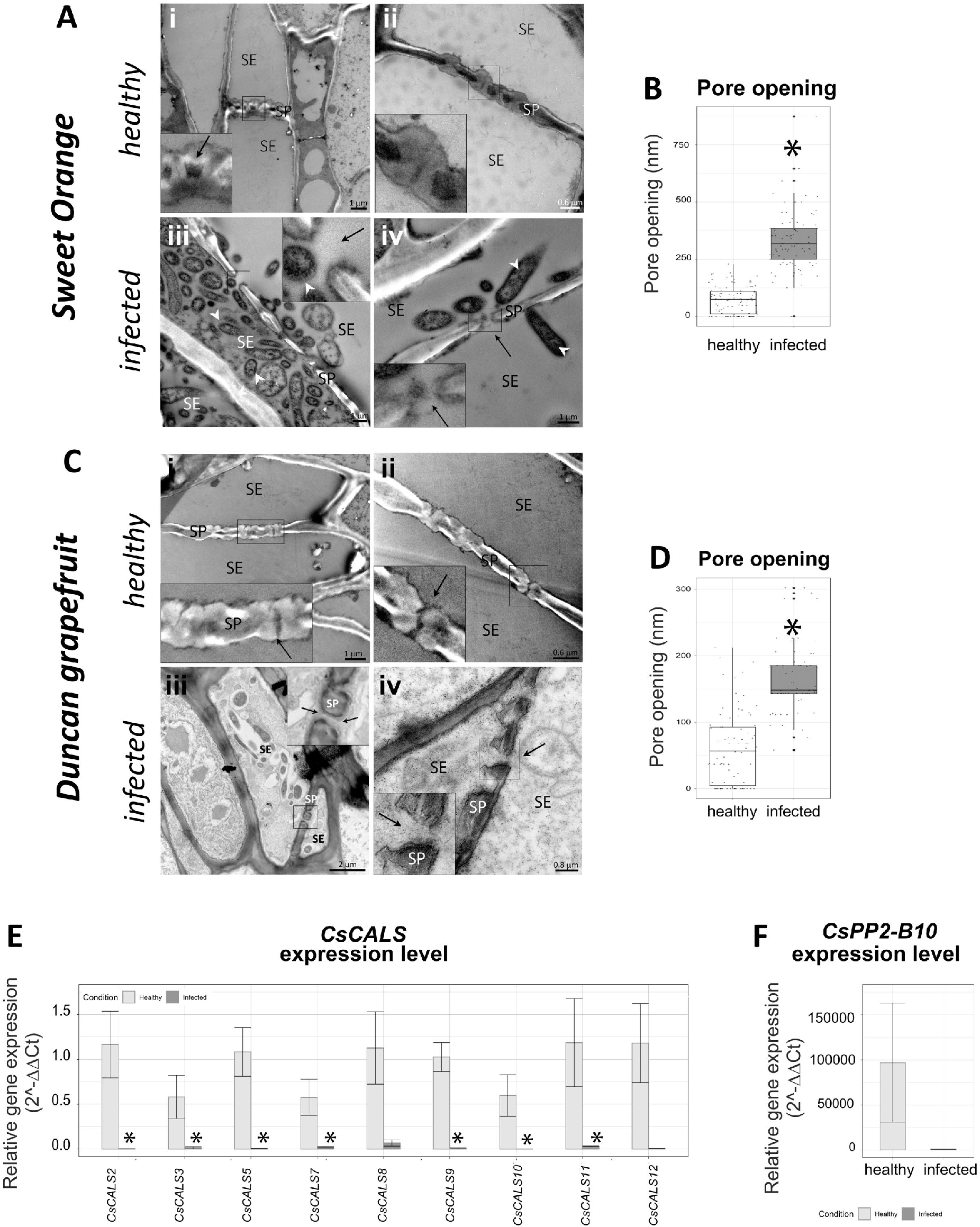
CLas inhibits phloem plugging in seed vasculature. A: Micrographs of healthy (i and ii) and infected (iii and iv) seed vasculatures of ‘Valencia’ sweet orange. SE= sieve element, SP = sieve plates, arrows= sieve pores, triangles= CLas; bar=1, 0.6, 1 and 1 μm respectively. B: Sieve plate pore size in sweet orange. Asterisks express significant differences among the means, with a *P* ≤ 0.05 (Student’s t test). C: Micrographs of healthy (i and ii) and infected (iii and iv) seed vasculatures of ‘Duncan’ grapefruit. Labels: SE= sieve element, SP = sieve plates, arrows= sieve pore channel and triangles= CLas; bar=1, 0.6, 2 and 0.8 μm respectively. D: Sieve plate pores of ‘Duncan’ grapefruit. Asterisks express significant differences among the means, with a *P* ≤ 0.05 (Student’s t test). E: Relative abundance of *CsCalS* gene transcripts in healthy and infected seed vasculature of ‘Duncan’ grapefruit. F: Relative abundance of CsPP2-B10 gene transcripts in healthy and infected seed vasculature of ‘Duncan’ grapefruit. Data are expressed as mean ±SE of 4 biological replicates. Differences among healthy and infected means were evaluated with Student’s t –test. Asterisks represent significant differences at *P* ≤ 0.05.

In infected seeds, the lumen of sieve elements was filled by bacteria and no callose layer was visible inside the pores. In the sweet orange seed phloem, the pore opening had an average diameter of 58.7 nm in healthy seeds and 126.2 nm in infected samples (Fig. 2B). In grapefruit, the average pore opening was 60.7 nm in healthy samples and 98.3 nm in infected seeds (Fig. 2D). No other differences were notable between the two groups of sample TEM images.

To further investigate the decreased accumulation of callose in infected seed vasculatures, the expression levels of callose synthase genes were measured. The analysis revealed a general downregulation for all the tested *CsCalS* genes in the infected seed vasculature (Fig. 2E). The expression levels of *CalS2, CalS3, CalS5, CalS7, CalS9, CalS10* and *CalS11* were significantly downregulated in the infected seed vasculatures compared to healthy ones. The expression of all *CalS* genes was reduced between 95 to 99.95% in infected samples compared to the healthy samples. The downregulation was statistically significant for all *CsCalS* except for *CalS8* and *CalS12*, where the *P* value was 0.059 and 0.055, respectively (Fig. 2E). The gene expression of the sweet orange *PP2-B10* was also reduced, albeit not significantly.

### Lack of callose allows for bacterial passage through the sieve pore

In the TEM images, in the absence of the callose layer bacteria were found inside the phloem pores, consistent with movement in the phloem, in both the sweet orange (Fig. 3A, B) and grapefruit sieve plates (Fig. 3C, D). We also observed the bacteria in different shapes close to the sieve pore(Fig. 3). During the crossing, these bacteria appeared to squeeze themselves through the pore, with the rod shape changing from rounded to elongated (Fig. 3). In all examples observed, the width of the elongated bacterial structure matched the size of the unplugged, callose-free pore.

**Figure 3:**
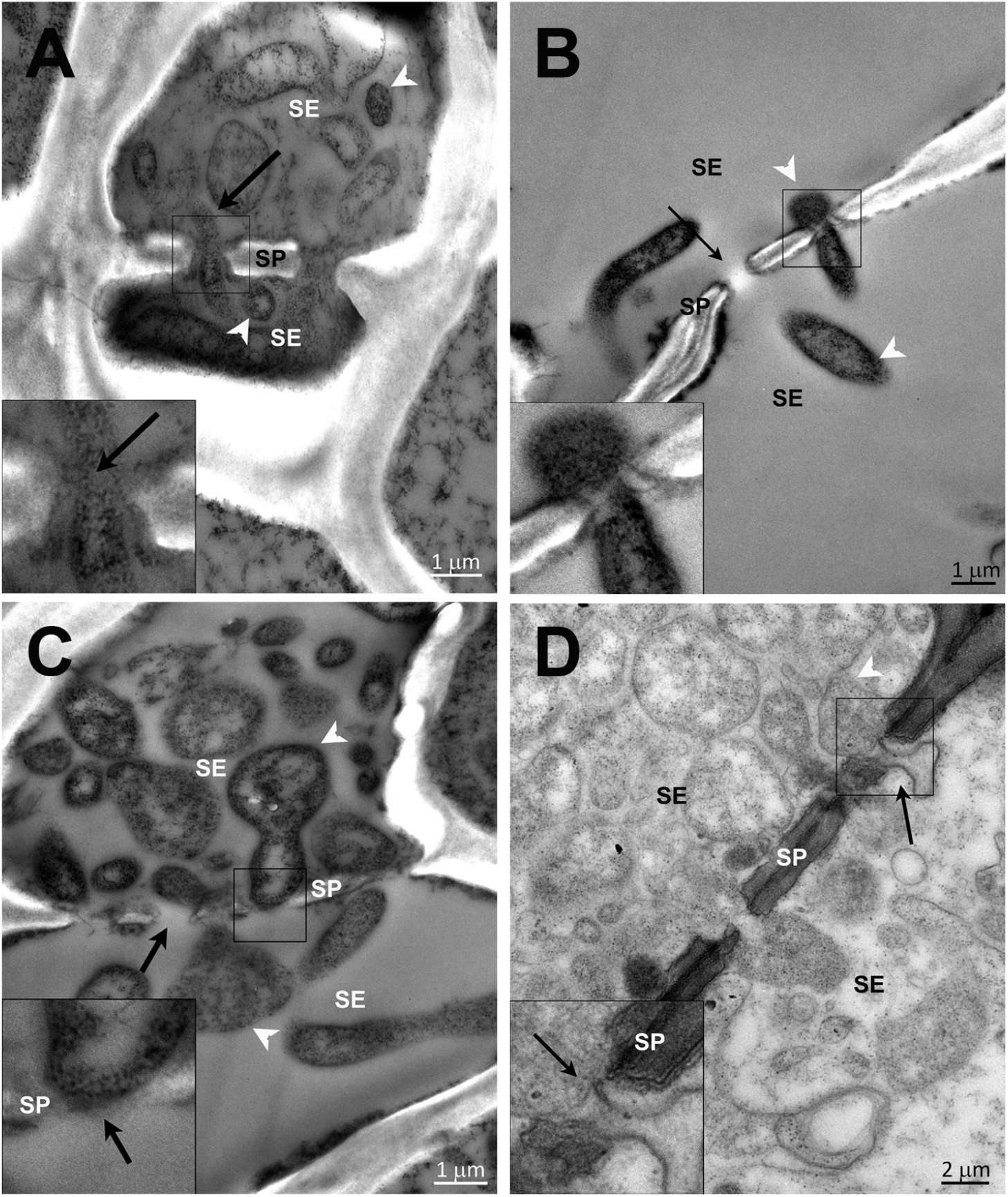
CLas passage through open sieve pores in seed vasculatures. Sieve pores of ‘Valencia’ sweet orange (**A-C**) and ‘Duncan’ grapefruit (**D**) with *C*Las. Arrows indicate the passage of *C*Las through the sieve pore without the callose layer. Bar= 1, 1, 1, and 2 μm respectively.

### H_2_O_2_ increases in infected young leaves, but decreases in infected seed vasculature

Previous research demonstrated that ROS was elevated in citrus leaves following *C*Las infection (Ma et al., 2022; Pitino et al., 2017). However, ROS has not been quantified in infected seed vasculatures, where our results indicated that bacteria are present in large numbers. In this study, leaves and seed vasculatures from healthy and infected samples were harvested and immediately stained in DAB solution, which turns red in the presence of H_2_O_2_. In the seed vasculatures, H_2_O_2_ levels were lower in infected samples compared to the healthy ones (Fig. 4A). Images of the vasculature samples were analyzed by a macro with FIJI to evaluate the intensity of the color. The average optical density (OD) value for healthy vasculature samples was 1.75 and the infected samples was 1.25 (Fig. 4B). Infected leaves had some red stained areas, indicating the presence of H_2_O_2_ (Fig. 4D). No red stain appeared in the healthy leaves. Healthy leaves showed an average optical density of 0.58 ± 0.17 and the infected leaves had an average optical density of 0.96 ± 0.25, confirming that H_2_O_2_ production increased in the *C*Las-infected leaves (Fig. 4E).

**Figure 4.**
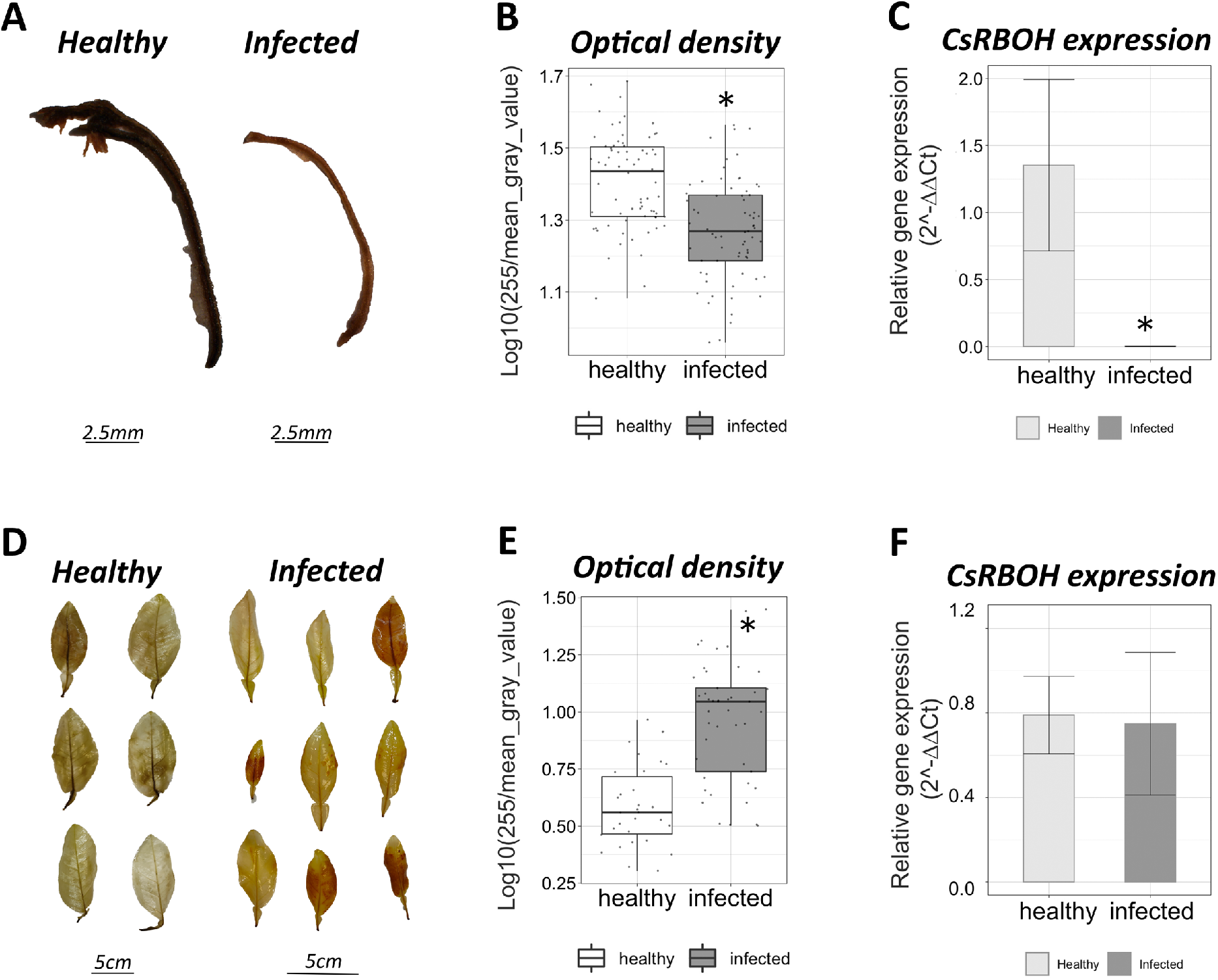
H_2_O_2_ production in seed vasculatures and leaves of ‘Duncan’ grapefruit. A: ‘Duncan’ grapefruit vasculatures stained with DAB (3,3’-diaminobenzidine) from healthy and infected seeds. Bar=2.5mm. B: Optical density of healthy and infected seed vasculatures. Asterisk expresses significant differences among the means (Student’s t-test, *P* ≤ 0.05). C: CsRBOH seed vasculature expression level in ‘Duncan’ grapefruit seed vasculatures expressed as mean ± SE. Asterisk denotes significant differences among the means (Student’s t-test, *P* ≤ 0.05). D: ‘Duncan’ grapefruit leaves stained with DAB (3,3’-diaminobenzidine) from healthy and infected plants. Bar=5cm. E: Optical density of healthy and infected leaves. Asterisk expresses significant differences among the means (Student’s t-test, *P* ≤ 0.05). F: *CsRBOH* seed vasculature expression level in ‘Duncan’ grapefruit seed vasculatures, expressed as mean ± SE. Asterisk expresses significant differences among the means (Student’s t-test, *P* ≤ 0.05).

To further analyze the DAB stain results, *CsRBOH* relative expression was analyzed in grapefruit leaves and seed vasculature. The RBOH protein is the provider of O_2_^-^ ions, a reactive oxygen species (Torres et al., 2006). In the infected seed vasculature, the *CsRBOH* gene was downregulated 1000-fold compared to healthy vasculatures (Fig. 4C). The average expression level was 1.353 ± 0.6300 in the healthy samples while in infected samples it was 0.001 ± 0.0003 (Fig. 4C). We did not observe a significant upregulation of *CsRBOH* gene expression in the leaves (Fig. 4F). The contrasting results from the leaves and seed vasculatures confirm that a strong inhibition of H_2_O_2_ production is caused by the presence of the pathogen inside the host cells.

### In infected young leaves, cells with *C*Las have less callose

Different varieties of HLB-susceptible citrus trees accumulate callose in the phloem of the leaves and stems (Welker et al, 2021; Granato et al., 2019; Achor et al., 2010). When observed under the TEM, healthy leaves show a layer of callose around the sieve pores. In HLB-infected trees, the sieve pores of the leaves are constricted by the callose, and this accumulation reduces the diameter of the pore by about 50% in both symptomatic and asymptomatic leaves (Achor et al., 2010), but *C*Las was not detectable in the sieve tubes in this study. Here, we carefully examined the sieve pores in leaf midrib sieve elements and identified *C*Las in a small fraction of the cells (Supplemental Table 1). On average, about 7.7% of the sieve element cells in the infected young leaves showed the presence of the *C*Las cells inside the lumen (Fig. 5A, C and Supplemental Table 1). We measured the sieve pore diameter in the sieve plates of cells with and without *C*Las (Fig. 5D). While both types of cells contained callose in the sieve pores, in the cells where *C*Las was present, there was a 2-fold increase in the pore diameter compared to the cells without *C*Las. These results suggest that when present, CLas in the sieve elements of leaves or seeds, either inhibits the deposition of callose or induces its removal.

**Figure 5.**
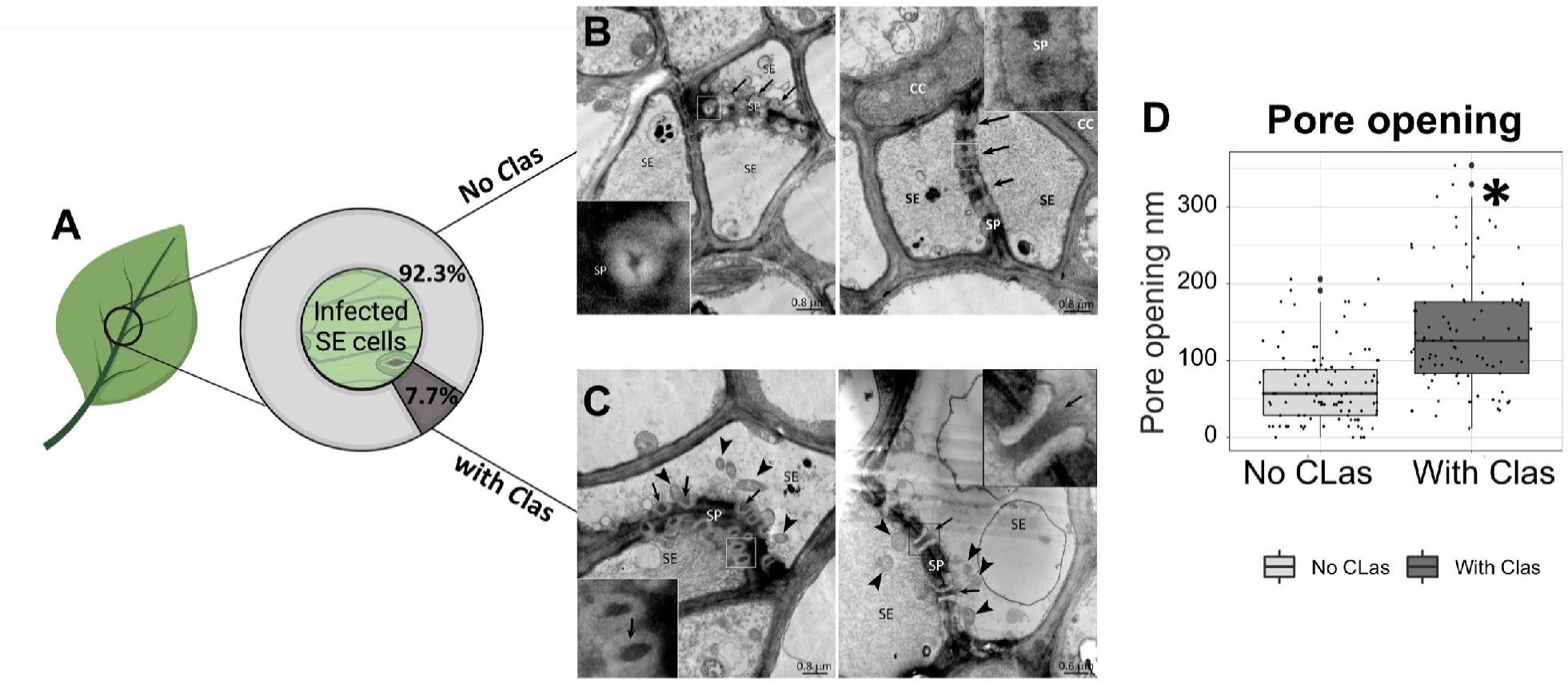
*C*Las reduce sieve pore callose in the leaves. A: Percentage of cells without *C*Las-(light grey) or with *C*Las (dark grey) in midrib of HLB infected plant. B-C: Micrograph of callose deposition in *CL*a-free (B) or *CLas*-containing (C) cells. Arrows= sieve pores, triangles= *C*las, SE= sieve elements, SP= sieve plates, CC= companion cells. D: Pore opening value in *CL*as-free and *CL*as-containing cells. The boxplot reports the pore opening in nm. Asterisk expresses significant differences among the means, with P ≤ 0.05 (Student’s t-test). Picture made with BioRender software (BioRender.com, 2022).

## DISCUSSION

In this study, we employed citrus seed vasculature, where *C*Las accumulates in large numbers, as a model tissue for the study of bacteria-plant interactions (Fig. 1). In the seed vasculature, the pattern of callose expression was the opposite of what was previously described in infected stems (Fig 2). Gene expression analysis confirmed the TEM observations and the imaging analysis of physical callose expression. Leaf sieve elements with living bacteria showed the same lack of callose accumulation as did the sieve elements in the seeds, indicating the response is not related to the type of tissue but rather to the presence of the pathogen (Fig. 5). In addition to reduced callose deposition (Fig. 2), H_2_O_2_ production was also reduced in *C*Las-containing cells (Fig. 4). This corresponded with a lower expression of the *CsCalS* and *CsRBOH* genes. Previously it was shown that callose and ROS metabolism increase in infected plants (Ma et al., 2022; Pitino et al., 2017; Granato et al., 2019), but these observations were made in the leaf tissue, where *C*Las levels are low. Our results suggest that these increases include both the activation by the host and inhibition by the pathogen, but because the pathogen is absent from most SE in the leaf, the net effect is an increase. The inhibition of the callose and ROS plant defense responses by *C*Las plays a key role for the efficient colonization of the host. Previous work has shown that the ability to bypass plant defenses in this manner would not be unusual among phytopathogens (da Cunha et al., 2007). In citrus, *Penicillium digitatum* can activate the catalase ROS scavenger machinery to limit the damage from H_2_O_2_ (Macarisin et al., 2007), and in the fruit this fungus can locally decrease the pH to enhance pathogenicity (Prusky et al., 2004).

A variety of possible methods by which *C*Las may suppress plant defenses have been described. This organism is known to control defense signaling through the bacterioferritin co-migratory protein (BCP) (Jain et al., 2021, 2019, 2018), through peroxidases (Jain et al., 2015), and through phytohormones (Li et al., 2017). The *C*Las genome encodes 86 proteins supposedly secreted by the Sec system (Xuelu Liu et al., 2019): ten of them appear to repress immune-mediated cell death and H_2_O_2_ production (Du et al., 2021). Two *C*Las proteins, m4405 and SDE15, were demonstrated to suppress cell death and downregulate *PR* genes (Zhang et al., 2020; Pang et al., 2020). *C*Las BCP (LasBCP) compromises the host systemic acquired resistance (SAR) response through an interaction with plant oxylipin metabolism and the lipopolysaccharides (LPS) on the bacterial surface (Jain et al., 2018; 2019; 2021). Surface LPS are known to provoke an immune response in the host plant. However, the expression of LasBCP in tobacco plants leads to a decrease of the LPS-triggered SAR (Jain et al., 2021). The LasBCP-modified tobacco plants also displayed a reduced deposition of callose (Jain et al., 2018), suggesting the generalized local inhibition of the response.

The local lack of callose deposition which was observed in this study could be due to the absence of an appropriate stimulus. Ca^2+^ is considered to be one of the stimuli for callose production (Brown and Lemmon, 2009). Decreases in calcium concentration were shown to occur following *C*Las infection, due to the increased activity of Calcium exchanger 7 (Xiaofei Liu et al., 2019). The local depletion of Ca^2+^ could impede the stimulus for the callose synthases. The reduction of the callose layer affects the size exclusion limit (SEL) of the sieve pores and consequently, the pathogen can easily migrate through the sieve plates from cell to cell (Fig. 3; Achor et al., 2020). Similarly, viruses also inhibit the deposition of callose layer around the sieve pore to facilitate movement through plasmodesmata (Guenoune-Gelbart et al., 2008; Rojas et al., 1997; Wang, 2021). Moreover, physiological studies have reported that both callose deposition and ROS production are enhanced by variation of calcium signature (Gilroy et al., 2016, 2014; Nedukha, 2015; Ogasawara et al., 2008).

*C*Las infection coincides with the upregulation of citrus *RBOH* and the downregulation of the ROS scavenger machinery, affecting the ascorbate peroxidase, catalase, and superoxide dismutase genes (Pitino et al., 2017). In the leaf tissue of infected plants, where the bacteria are not present, there is an increased level of H_2_O_2_. Information gleaned from the *C*Las genome sheds light on the cause of reduced H_2_O_2_ in the tissues where the *C*Las bacteria are highly numerous. Two *C*Las prophages encode for putative peroxidase: SC2_gp095 and the glutathione peroxidase SC1_gp100 (Jain et al., 2015; Zhang et al., 2011). SC2_gp095 can degrade H_2_O_2_, while SC2_gp095 suppresses the activation of RBOH to avoid the toxicity of ROS thus allowing the bacteria to survive (Jain et al., 2015). Results from this work are summarized in Figure 6. Healthy plant sieve elements have a layer of callose around the sieve pore and a physiological level of ROS, salicylic acid (Nehela et al., 2018), and Ca^2+^ (Hijaz et al., 2016). Sieve elements that do not contain bacteria display the opposite pattern if the phloem, as a whole, is infected with HLB. In these cells, there is an increase of Ca^2+^ and SA (Xiaofei Liu et al., 2019; Nehela et al., 2018), leading to an increase of callose and ROS (Fig. 4). The intense immune response from the citrus host results in a reduction of photosynthate transport ability and the eventual destruction of important phloem and photosynthetic structures (Achor et al., 2010; Folimonova and Achor, 2010; Granato et al., 2019; Pitino et al., 2017; Welker et al., 2021). In cells where *C*Las is present, callose and ROS are reduced, and the relevant metabolic pathways are downregulated. Thus, the bacteria can survive and move, and profit from the host metabolism despite the plant’s strenuous opposition.

**Figure 6.**
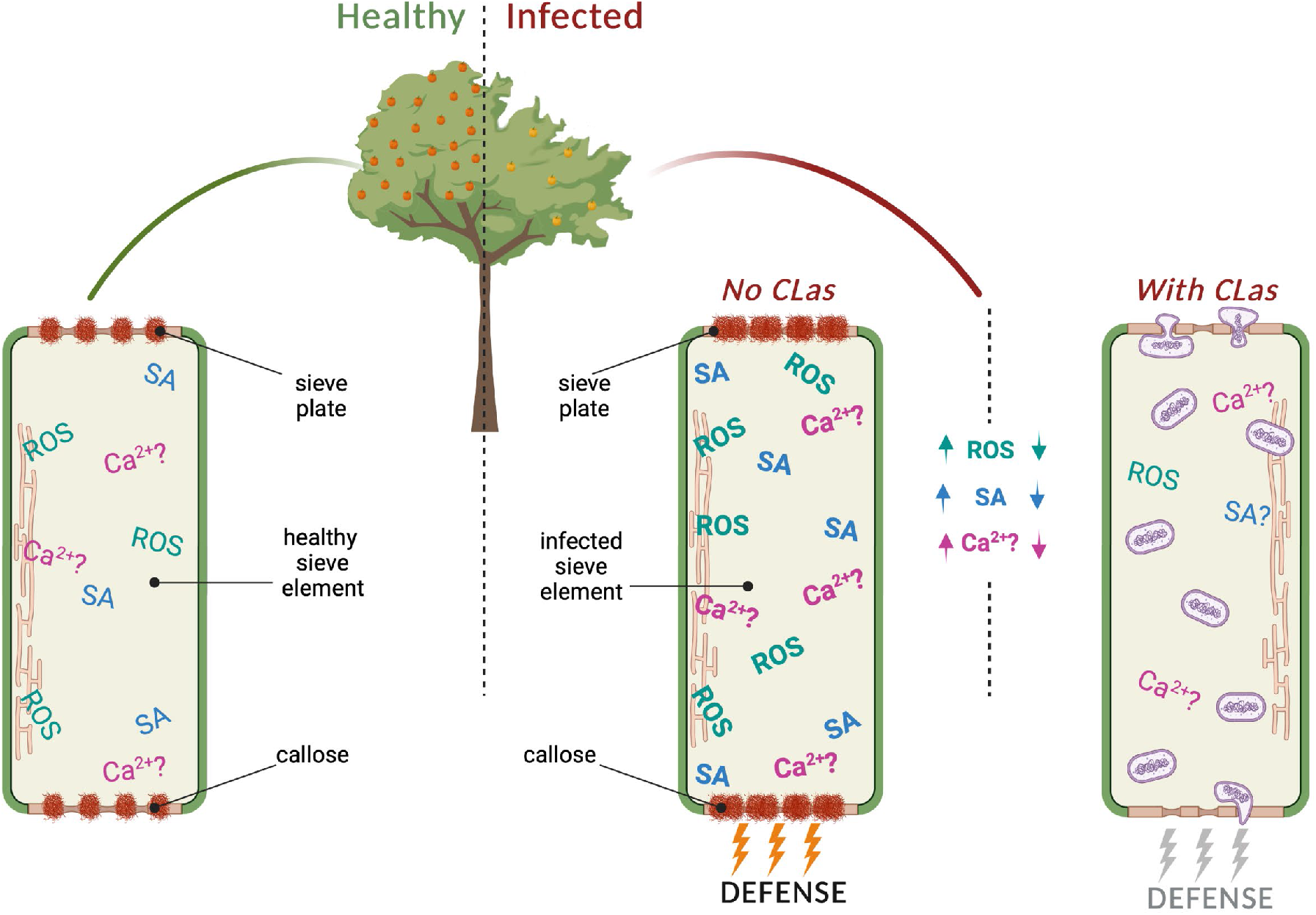
Model for CLas-phloem interaction in HLB-infected trees. In healthy sieve elements, a physiological level of ROS (reactive oxygen species), SA (salicylic acid) and Ca^2+^ is present in the phloem sap. Around the sieve pores, a normal physiological layer of callose ensures proper transport of substances through the phloem. In infected sieve element cells without *C*Las, the ROS concentration and the SA increase. Ca^2+^ may increase as well. Callose completely occludes the sieve pores. Defense responses in the cell are activated due to the perception of the pathogen by the plant. In CLas-containing sieve element cells, ROS concentration decreases. The concentration of Ca^2+^ and SA probably decrease as well. Sieve pore callose is completely absent, allowing the movement of the bacteria. ROS= reactive oxygen species, SA=salicylic acid, Ca^2+^=calcium ions. Picture made with BioRender software (BioRender.com, 2022).

HLB disease symptoms appear to result from a failed strategy on the part of the citrus host, where the cost of defense outweighs the damage done by the bacterial pathogen itself. Analysis of the *C*Las genome reveals an organism with largely defensive capabilities, which makes sense for an obligate parasite of a living host (Duan et al., 2009). Blocking the attempts of *C*Las to bypass the plant defenses can provide one strategy for eliminating the bacteria and the establishment of resistant varieties. Paradoxically, the bacterial activities that counteract the host defenses may also alleviate some disease symptoms. In some cases, plants which have modifications that result in less callose expression have shown better resistance to powdery mildew (Jacobs et al., 2003; Nishimura et al., 2003). This phenomenon has not been greatly studied in the context of bacterial pathogens which inhabit the phloem. Considering this information, a “non-confrontational” strategy for the development of HLB-resistant citrus might also be considered. Hypothetically, citrus stock which naturally exhibits a low callose and ROS response to the presence of *C*Las might also remain free from the symptoms of HLB (Curtolo et al., 2020; Deng et al., 2019). Future work could focus on screening existing citrus varieties based on these criteria to see if they survive well despite the pathogen presence. This strategy might also be examined in other plant-pathogen interactions which involve bacteria inhabiting the vascular system.

## Conclusion

We show that the reduction of phloem sieve plate pore callose and inhibition of plant ROS production by *C*Las are two key aspects for the establishment of HLB disease in the host. The local inhibition of ROS enhances pathogen survivability while the reduction of the callose allows systemic colonization of the plant. This information can be used to better understand the nature of plant diseases which involve obligate parasites of the phloem.

## Materials and methods

### Sample collection

Fruits and flush leaves of *Candidatus* Liberibacter asiaticus (*C*Las)-infected ‘Valencia’ sweet orange (*Citrus sinensis* L.) and ‘Duncan’ grapefruit (*Citrus* x *paradisi*) (DG) were collected simultaneously in experimental fields in Polk County (July 2020) and Collier County (July 2021), Florida, USA at the fully symptomatic stage. Healthy fruits and leaves belonging to the same varieties were collected at the same time from plants grown under protective screens. Healthy samples were used as a control group.

Seeds were collected from fruit stored at 4°C until seed vasculature extraction. The testa and the tegmen of the seed were removed with forceps to expose the vascular tissue of the embryo. The vasculatures were carefully removed from the apex at the point of junction with the embryo (Supplementary Figure 1). The extracted vasculatures were used either immediately (microscopy analysis), stored at 4°C for staining, or stored at −20°C for molecular analysis.

### Electron microscopy and imaging analysis

Electron microscopy analysis was performed as previously described (Folimonova and Anchor, 2010, and Anchor 2020), using a standard fixation procedure as follows. Midrib samples (0.5mm of length) and seed vasculature samples (whole vasculature), were collected from 3 different plants. Samples were fixed with 3% (v/v) glutaraldehyde in 0.1 M of potassium phosphate buffer at pH 7.2 for 4 h at room temperature, washed in phosphate buffer, then postfixed in 2% osmium tetroxide (w/v) in the same buffer for 4 h at room temperature. The samples were further washed in the phosphate buffer, dehydrated in a 10% acetone (v/v) series (10 min per step), and infiltrated and embedded in Spurr’s resin over 3 d. Sections (100-nm) were mounted on 200-mesh formvar-coated copper grids, stained with 2% aq uranyl acetate (w/v) and Reynold’s lead citrate. Ultrathin sections of the samples were observed through a Morgagni 268 transmission electron microscope (TEM). Pictures obtained from the TEM observations were analyzed with FIJI. For each sieve plate, the opening of the pores was measured, and analyzed as reported below.

### Confirming the accumulation of Clas in the seed vasculature

The accumulation of the bacteria in the seed vasculature was assessed with fluorescent in situ hybridization following a previously reported protocol (Ghanim et al., 2009) and adapted for use with plant tissue (Hilf et al., 2013). Samples were visualized with a Leica SP8 laser-scanning confocal microscope (Leica Microsystems Inc., Buffalo Grove, IL).

### H_2_O_2_ concentration assay

Samples of leaves were collected from the field and immediately placed in a solution of 1mg/ml 3,3’-diaminobenzidine (Sigma-Aldrich) (DAB) in water, pH 3.8, prepared as previously reported (Daudi and O’Brien, 2012; Kumar et al., 2014). Seed vasculatures collected from the fruits, stored at 4°C, were placed in DAB solution. The staining protocols were performed as follows: overnight staining in DAB on a shaker at 50 rpm. The following day destain for 20 minutes (for leaf samples) or 10 minutes (for seed vasculature samples) in a bleaching solution with the composition of ethanol:glycerol:acetic acid in a 3:1:1 ratio at 92°C (Daudi and O’Brien, 2012; Kumar et al., 2014). After the initial bleaching, the solution was replaced with fresh bleaching solution and the samples were left at room temperature for 30 minutes before the observation. Leaf samples were collected from 10 plants (5 healthy and 5 infected) and at least 6 leaves selected randomly in each plant were observed. For seed vasculature analysis, seed vasculatures were extracted from 3 fruits chosen randomly from each of 10 plants (5 healthy and 5 infected). For each fruit at least 10 seed vasculatures were observed, and pictures were acquired with a stereo microscope Leica KL300 on a white background (Leica Microsystem 2021) and using the LASX software (Leica Application Suite, Leica microsystem 2021). For each picture the same exposure time, light and white balance conditions were applied.

Leaf pictures were analyzed with FIJI software (Schindelin et al., 2012). On each picture, color deconvolution was performed with the H DAB algorithm (Crowe and Yue, 2019). The channel corresponding to the DAB color was separately saved and used for the analysis. For each leaf the mean gray value (intensity of the light) of the whole lamina was analyzed, extracting the lamina from the background.

Vasculature tissue was analyzed with the workflow as described above. For each vasculature, 5 regions of interest (ROIs), each 576 pixels squared in size, were chosen randomly to avoid readings affected by the size of the vasculature. For each ROI, the mean gray value was recorded.

The mean gray value was transformed to optical density value with the following formula: log(max intensity / mean intensity) in which the maximum intensity for 8-bit pictures has a value of 255 (Nguyen and Nguyen, 2013).

### Gene expression analysis

Total RNA extraction on Duncan seed vasculatures and leaf midribs were performed using Trizol reagent (Invitrogen). Quantitative PCR was performed as previously described (Achor et al., 2010), starting from roughly 100 mg of fresh tissue. RNA reverse transcription was carried out using a High-Capacity cDNA Reverse Transcription kit (Applied Biosystems), following the manufacturer instructions starting from 500 ng of RNA. To determine difference in modulation of the genes, RT-PCR was performed using SYBR Green FastMix (Quantabio, Beverly, MA, USA) starting from 300 ng of cDNA in a total reaction volume of 15 μL with a concentration of each primer of 400 nM. All gene primers used for RT-PCR are listed in Table 1. Gene expression was compared to the citrus GAPDH reference gene, and analysis was performed using the 2^-ΔΔCt^ method (Livak and Schmittgen, 2001).

**Table 1:**
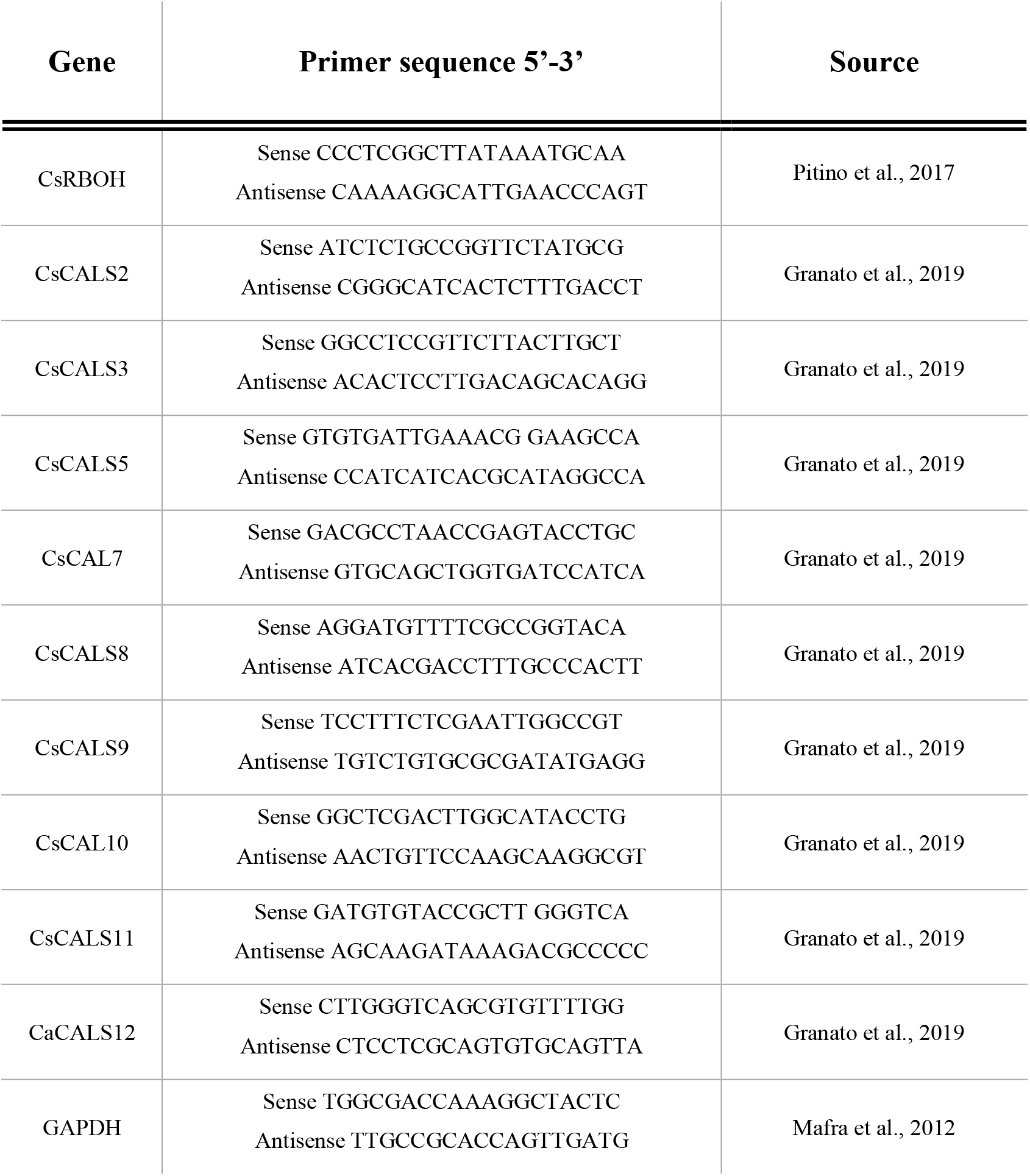
List of the genes analyzed in this study and relative primers used to amplify them.

### Data analysis

Statistical analyses were performed using R with Rstudio software Version 1.1.456 (RStudio Team (2020). RStudio: Integrated Development for R. RStudio, PBC, Boston, MA). Conformity to the normal distribution and homogeneity of variances were checked with Shapiro-Wilk’s test and Bartlett’s test respectively. Where necessary, data were normalized with a Box-Cox transformation. For each analysis, Student’s t-test was used to determine significant differences among the treatment group means (healthy or infected) with *p* < 0.05.

## Acknowledgement

We thank Dr. Vladimir Orbovic, Dr. Arnold Scumann, Dr. Fernando Alferez and Laura Waldo (University of Florida) for providing us grapefruit and sweet orange fruits. This research was funded by the National Institute of Food and Agriculture, grant number 2020-70029-33197 to Amit Levy and Robert Turgeon.

## Author contributions

AL conceived the project. CB CW and DT collected the samples. CB performed the ROS analyses, the statistical analysis. CB and DT carried out the gene expression analysis. DA and CW embedded the samples and visualized by TEM. YAA performed the FISH analysis. AL and CB performed the imaging analyses. CB, AL, RT and SW wrote the manuscript. All the coauthors provided critical comments for the text of the manuscript.

